# SECRET AGENT O-GlcNAc Modifies GIGANTEA: Methods for Mapping of O-Linked β-N-Acetylglucosamine Modification Sites Using Mass Spectrometry

**DOI:** 10.1101/2023.04.13.536733

**Authors:** Young-Cheon Kim, Lynn M. Hartweck, Neil E. Olszewski

## Abstract

*Arabidopsis thaliana* has two glycosyl transferases, SPINDLY (SPY) and SECRET AGENT (SEC), that modify nuclear and cytosolic protein with O-linked fucose and N-acetylglucosamine (GlcNAc), respectively. SPY interacts physically and genetically with GIGANTEA (GI). Previously, we reported that SEC substrates are O-GlcNAc modified when they are co-expressed in *E. coli*. By analyzing overlapping sub-fragments of GI, we found a region that was modified by SEC. Mutational mapping of the modified region was then performed. Modification was undetectable when threonine 829 was mutated to alanine (T829A) while the T834A and T837A mutations reduced modification suggesting that T829 was the primary or only modification site. Mapping using several enrichment and mass spectrometry methods all detected only modification of T829.

## INTRODUCTION

Post translational modification of serine or threonine residues with O-Linked β-N-acetylglucosamine (O-GlcNAc) regulates the activity of nuclear and cytosolic proteins (Ma et al., 2022). *Arabidopsis thaliana* has one O-GlcNAc transferase, SECRET AGENT (SEC) (Hartweck et al., 2002). Arabidopsis also has an O-fucose transferase, SPINDLY (SPY) (Zentella et al., 2017). Both enzymes modify serine and threonine of nuclear and cytosolic proteins. *sec* mutants have subtle plant defects while *spy* mutant have more and stronger defects (Hartweck et al., 2006; Hartweck et al., 2002; Sun, 2021; Zentella et al., 2016). *sec spy* double mutants are embryo lethal suggesting that SEC and SPY have overlapping functions (Hartweck et al., 2002). Recent surveys have identified numerous O-GlcNAc and O-fucose modified plant protein (Bi et al., 2023; Li et al., 2023; Wu et al., 2022; Xu et al., 2017; Zentella et al., 2023). Consistent with SEC and SPY having overlapping functions, O-GlcNAc and O-fucose modifications can occur at the same position (Bi et al., 2023; Xu et al., 2017).

Previously, we showed that GIGANTEA (GI), a protein that regulates many processes including circadian clock, flowering time, and light signaling (Brandoli et al., 2020; Krahmer et al., 2019; Mishra & Panigrahi, 2015), interacts physically and genetically with SPY (Tseng et al., 2004). We also have shown that when co-expressed in *E. coli* SEC modifies GI (Kim et al., 2013). In this study we used deletion analysis, site directed mutagenesis and mass spectrometry to map where SEC modifies GI. We also investigated several methods for enriching and mapping O-GlcNAc modification.

## MATERIALS AND METHODS

### GI expression constructs

To make the *E. coli* expression constructs, fragments of the GI protein coding region were amplified by PCR using the primers listed in Supplemental Table 1 and cloned between the BamHI and NotI sites of pET32a such that GI is produced with N-terminal S- and His-tags. To make the CT5 construct the region encoding GI amino acids 789-893 was cloned between the NcoI and XhoI sites of pET32a. Site directed mutagenesis was performed using the QuikChange Site-directed Mutagenesis Kit following the instructions of the manufacturer (Stratagene; La Jolla, CA) using the primers listed in Supplemental Table 1.

### Detection of GlcNAcylated proteins on protein blots

SEC and different portions of GI were co-expressed in the *E. coli* strain BL21-AI as described previous (Scott et al., 2006). Cells were lysed, total proteins were resolved by SDS PAGE, blotted to PVDF membrane and the blots were probed to detect GlcNAc modified proteins using a radioactive galactosyl transferase assay that caps GlcNAc with ^3^H-gal as described previously (Scott et al., 2006).

### Recombinant CT5 purification and enrichment of modified CT5 peptides using RCA I lectin and avidin

O-GlcNAc modified CT5 was prepared and purified as described previously (Kim et al., 2011). Details of RCA I affinity chromatography procedures are described in (Kim et al., 2011). Before lectin and avidin gel enrichment, the CT5 proteins were digested with trypsin (40:1) or Lys-C (100:1) overnight at 37 °C, and then desalted using Sep-Pak C_18_ cartridge (Waters) and dried using a SpeedVac. Biotinylation, the addition of biotin to terminal GlcNAc modifications, proceeds in two steps. Enzymatic labeling was done with the Click O-GlcNAc enzymatic labeling system (Invitrogen) following the manufacturer’s instructions. After the labeling reaction, the sample was desalted using Sep-Pak C_18_ cartridge, and then dried in a SpeedVac. Biotin labeling was done with Click-iT protein analysis detection kit following the manufacturer’s instruction. Biotin labeled peptide was desalted using SCX spin column (Nest Group), and then purified by monomeric avidin gel following the manufacturer’s protocol (Pierce). Degalactosylation was done with β1-4 galactosidase following the manufacturer’s instruction (New England Biolabs), and desalted and dried using Sep-Pak C_18_ cartridge and SpeedVac. For the β-elimination reaction, desalted and dried peptides were resuspended in 1.5 % Triethylamine and 0.15 % NaOH, and then incubated at 52 °C for 1.5 hr. After β-elimination, the sample was desalted using Sep-Pak C_18_ and dried in a SpeedVac prior to the next procedure or MS analysis.

### Matrix-assisted laser desorption ionization time-of-flight (MALDI-TOF) analysis

Peptide samples were purified with C_18_ zip-tip (Millipore). 1.2 μl of elution were mixed on the target with matrix (10 mg/ml α-cyano-4-hydroxycinnamic acid in 50 % acetonitrile, 0.1 % trifluoroacetic acid) and analyzed in reflector mode on a matrix-assisted laser desorption ionization time-of-flight MS (MALDI-TOF) Bruker Reflex III (Bruker Daltonics). Processing of the spectra and data analysis was performed with Bruker Daltonics XTOF 3.1.

### Nanoflow LC – MS/MS and analysis of MS/MS data

Reversed-phase nano HPLC online with electrospray ionization mass spectrometry using a LTQ-OrbitrapXL equipped with electron transfer dissociation (ETD) and collision induced dissociation (CID) sources (ThermoScientific) was performed as described previously (Bandhakavi et al., 2009; Kim et al., 2013).

## RESULTS

### Mapping where SEC modifies GI by deletion and mutational analysis

Since we were unable to co-express full-length GI protein with SEC in *E. coli,* smaller segments that collectively span GI were co-expressed and analyzed for O-GlcNAc modification (Figure 1). All fragments that contained amino acids 828 to 840 were modified suggesting this region is modified.

**Figure 1.**
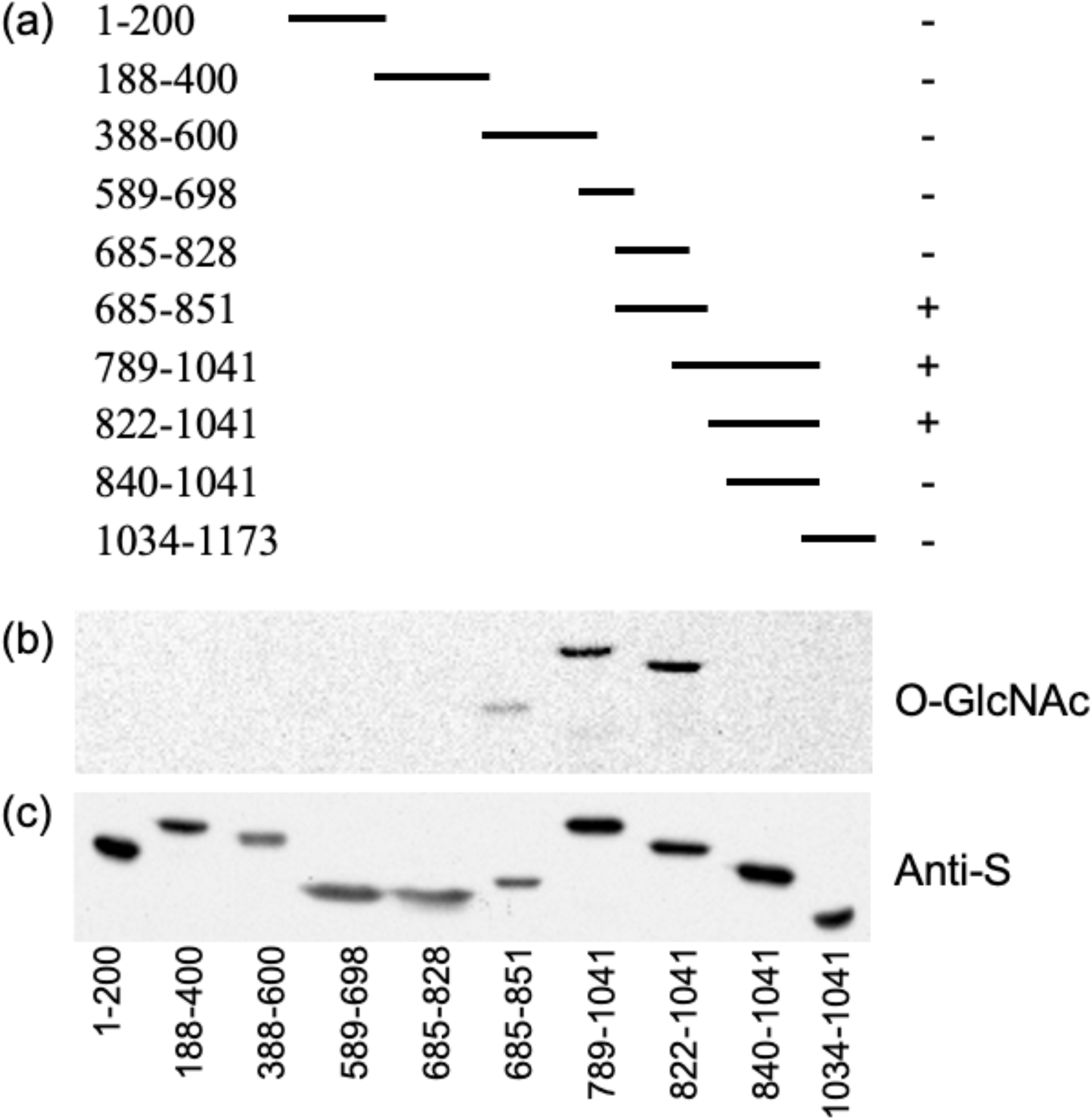
SEC modifies the region encompassing amino acids 828 to 840 of GI. (a) Map of the segments of GI that were co-expressed with SEC. + indicates the protein was modified and – indicates it was not. (b) GlcNAc modification of blotted proteins was detected using a galactosyl transferase assay (O-GlcNAc). (c) Duplicate blot probed with anti-S antibody to confirm the expression of GI (Anti-S).

A construct called CT5 which has GI amino acids 789 to 893, expressed well and was highly modified (Supplemental Figure 1a). Therefore, this construct was used for further mapping studies. The serine (S) and threonine (T) residues of the region spanning amino acids 825 to 840 were individually mutated to alanine (A) and the mutant proteins were tested to determine if they could be modified by SEC (Figure 2). T834A and T837A mutations reduced modification and modification of the T829A mutant was not detectable. Since mutations can affect protein structure or may affect the interaction with SEC it is not clear if the effects of the mutations are due to disruption of a modification site or an indirect effect on modification at another site. Therefore, mass spectrometry (MS) mapping was employed to map the modification site(s).

**Figure 2.**
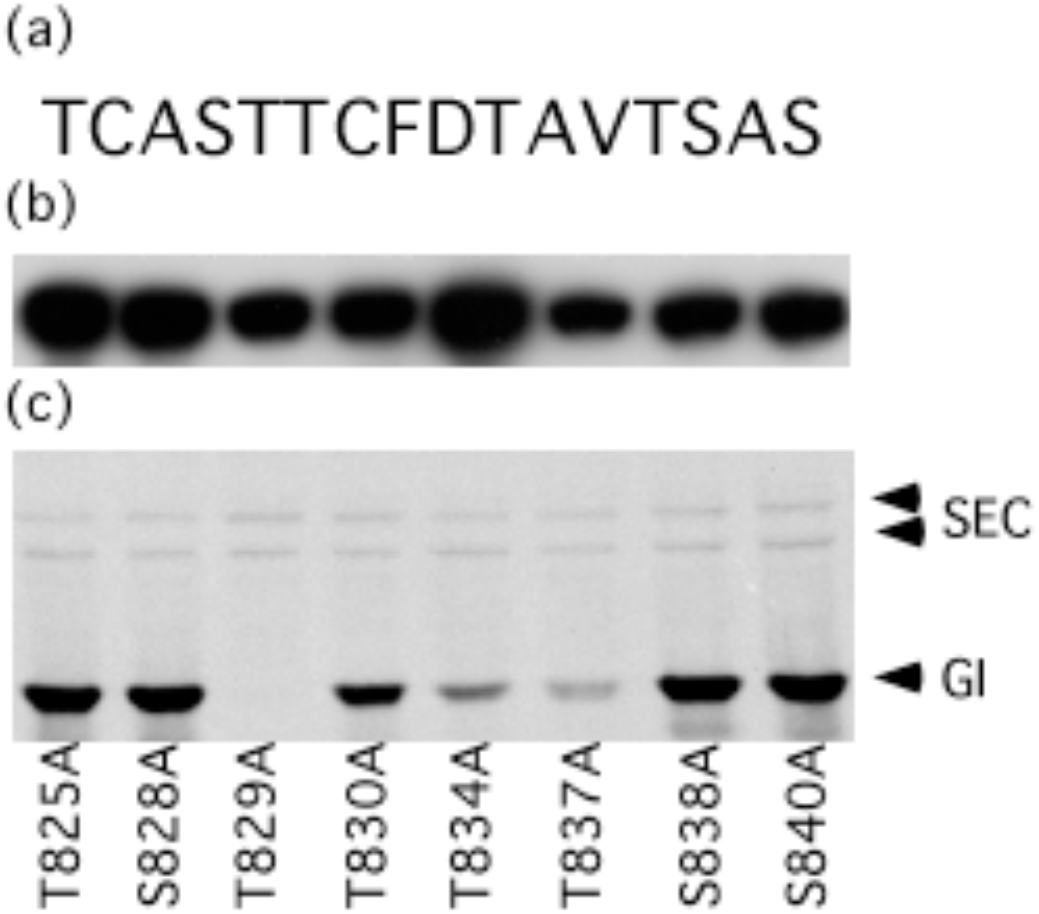
Mutational mapping of GI O-GlcNAc modification sites. (a) Amino acid sequence of GI 825-840 region. (b) Fluorograph detecting O-GlcNAc modification of mutant GI proteins. (c) Western blot using anti-S antibody.

### Strategies for enrichment of O-GlcNAc modified peptides

We used several different methods to enrich modified CT5 peptides and map the modifications (Figure 3). In the first method, O-GlcNAc was capped with galactose creating the disaccharide N-acetyl-D-lactosamine (LacNAc) and the modified peptides were enriched by *Ricinus communis agglutinin* I (RCA I) lectin affinity chromatography (Figure 3a). In the second method, modified sites were labeled with GalNAz and biotinylated using click chemistry. The biotinylated peptides were purified using avidin gel, and then the label including GlcNAc was removed in a β-elimination reaction producing dehydrated serine or threonine at the modified site. Since N-linked modifications are β-elimination-refractory, a second avidin affinity purification step was performed, to remove any GlcNAc modified peptides (Figure 3b). The third method is a combination of the first two methods (Figure 3c).

**Figure 3.**
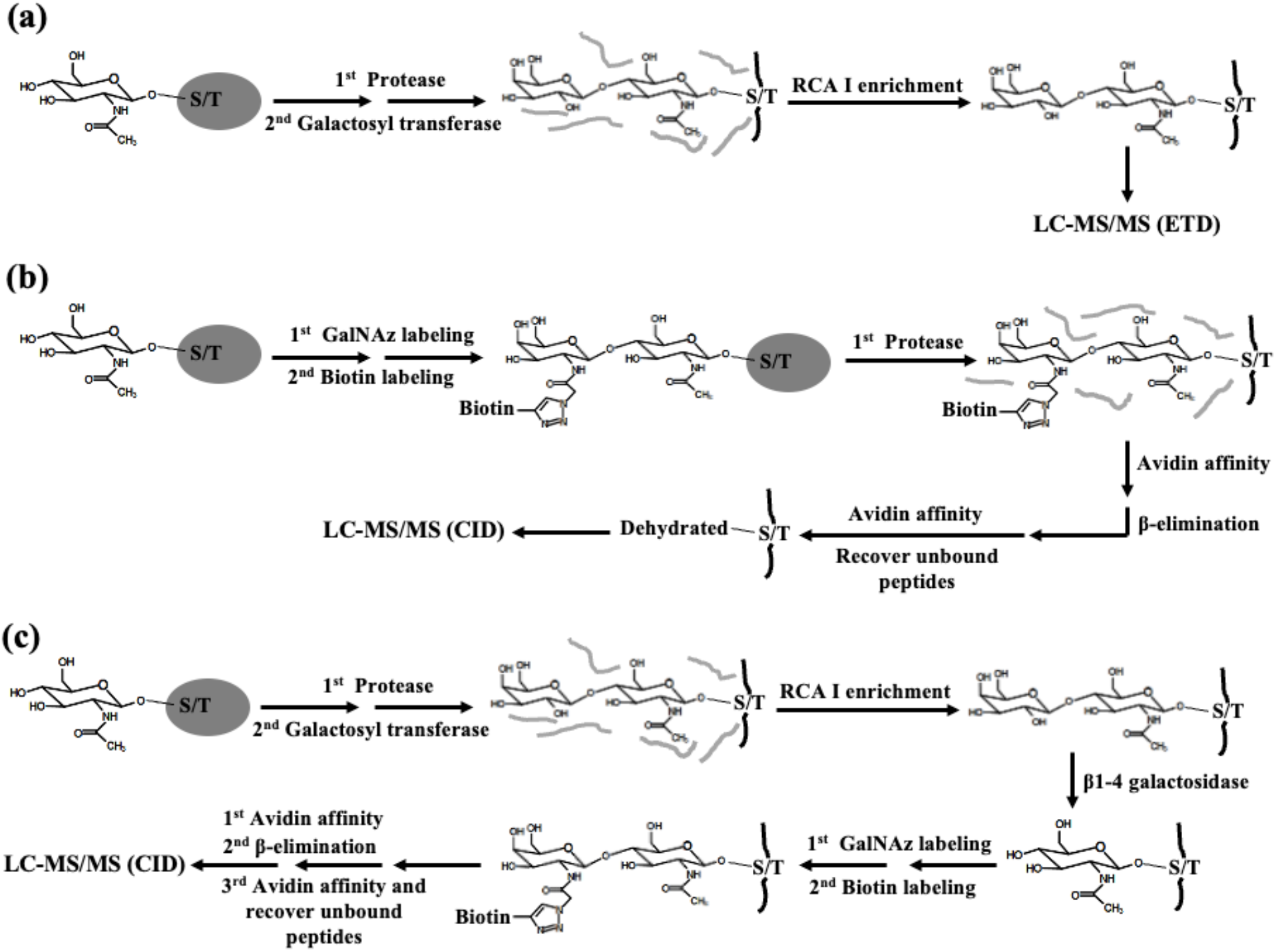
Strategies for enrichment and analysis of O-GlcNAc modified peptides. (a) O-GlcNAc modified peptides are capped with galactose using β1-4 galactosyltransferase. The modified peptides are enriched by RCA I affinity chromatography and the modifications are mapped using ETD mass spectrometry. (b) Modified proteins are labeled with GalNAz and biotin using Click enzymatic/chemical reaction. Trypsinized peptides are enriched by avidin affinity, and enriched peptides are subjected to β-elimination. Unbound peptides (flow through) are recovered from a second avidin affinity purification procedure and analyzed by CID mass spectrometry. (c) Modified peptides capped galactose are enriched by RCA I affinity chromatography and degalacosylated by β1-4 galactosidase. GlcNAc residues are labeled GalNAz and biotin and enriched by avidin affinity. Enriched peptides are subjected to β-elimination and unbound peptides are recovered from an avidin affinity purification procedure and analyzed by CID mass spectrometry.

The modified sites of the peptides purified using the first method were mapped using electron transfer dissociation (ETD) mass spectrometry which retains GlcNAc moieties. In contrast, O-GlcNAc modifications are unstable during collision induced dissociation (CID) mass spectrometry. Therefore in the second and third methods enriched peptides were subjected to β-elimination, which removes O-modifications and dehydrates the modified amino acid. The resulting dehydrated amino acids were mapped by CID mass spectrometry.

### Enrichment of O-GlcNAc modified peptides using lectin and avidin affinity chromatography

Matrix-assisted laser desorption ionization time-of-flight (MALDI-TOF) of trypsin-digested CT5 identified unmodified peptide (*m/z* 2209) and a peptide with a single O-GlcNAc (203 Da) fragment (*m/z* 2412) (Supplemental Figure 1a) indicating that CT5 is only partially modified when co-expressed in *E. coli* with SEC. MALDI-TOF analysis of peptides enriched by RCA I lectin affinity chromatography demonstrated successful enrichment of modified peptides produced with either trypsin (*m/z* 2573) or Lys-C digestion (*m/z* 3736) (Supplemental Figure 1b).

O-GlcNAc modified peptides that had been labeled by GalNAz and biotinylated using click chemistry (*m/z* 3183) were also successfully enriched using avidin gel (Supplemental Figure 2). When enriching peptides by combining the RCA I lectin and avidin gel approaches, the peptides from the RCA I enrichment must be treated with β1-4 galactosidase to remove galactose prior to GalNAz labeling. When galactose removal was monitored by MALDI-TOF, it was found to be nearly complete after 15 hrs (Supplemental Figure 3).

### Mapping of O-GlcNAc modification sites using ETD and CID mass spectrometry

Peptide enriched by RCA I after Lys-C digestion was mapped using ETD mass spectrometry, a quintuple-charged precursor ion (751.3275 mass-to-charge (*m/z))* with the sequence QENTCASTTCFDTAVTSASRTEMNPRGNHK had c_8_ and z23 ions with a 365 Da (LacNAc) mass increase which supports O-GlcNAc modification of T41 of CT5, which corresponds to T829 of full-length GI (Figure 4a). When β-eliminated peptide that had been purified by avidin affinity was analyzed by CID, a double-charged precursor ion 1003.9561 *m/z* corresponding to dehydrated QENTCASTTCFDTAVTSASR peptide was detected and b_8_ and y_13_ ion indicated that T41 had been dehydrated (Figure 4b). T41 of β-eliminated peptides enriched by RCA I lectin and avidin affinity was also modified at this position (Figure 4c). Modification at other positions was not detected with any of the methods.

**Figure 4.**
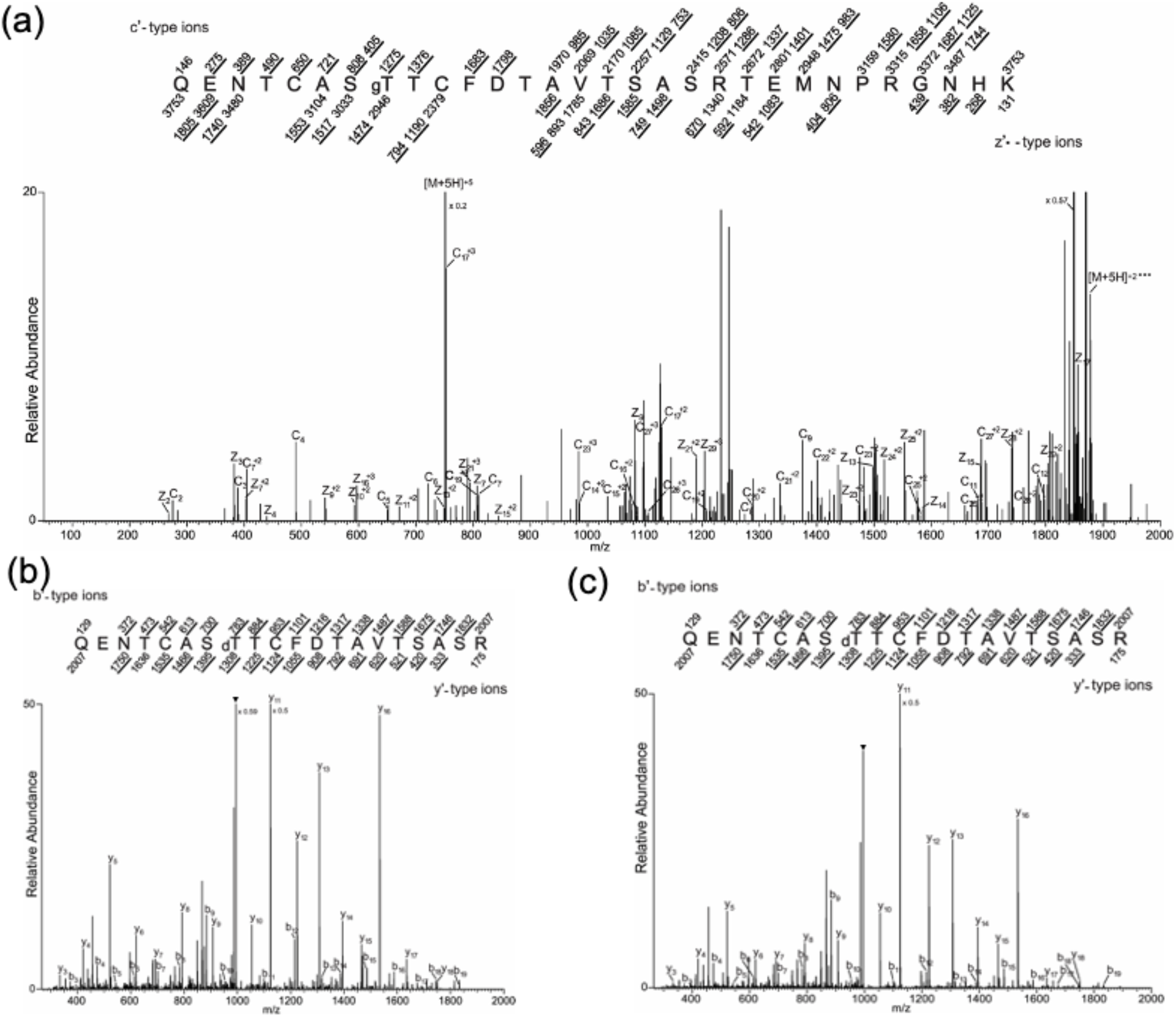
Mapping of CT5 modification site by mass spectrometry. (a) CT5 proteins were digested with Lys-C, GlcNAc was capped with galactose and modified peptides were enriched by RCA I affinity chromatography (Figure 1A). The ETD MS/MS spectrum recorded on [M+5H]^+5^ ions (*m/z* 751.3275) from LacNAc (365.1322) modified CT5 peptide QENTCASTgTCFDTAVTSASRTEMNPRGNHK. Predicted c’ and z’•-type ions are listed above and below the peptide sequence, respectively. Singly and doubly charged fragment ions are listed as monoisotopic masses. Ions observed and labeled in the spectrum are underlined. Modified residues are preceded by “g” to signify modification by a single LacNAc moiety. (b) and (c) Trypsinized CT5 peptides were prepared by Figure 1a and c method, respectively. The CID MS/MS spectrum was recorded on [M+2H]^+2^ ions (*m/z* 1003.9561 and 1003.9565) corresponding to the peptide QENTCASTdTCFDTAVTSASR dehydrated at T41. Predicted b’ and y’-type ions are listed above and below the peptide sequence, respectively. Singly charged fragment ions are listed as monoisotopic masses, respectively. Ions observed and labeled in the spectrum are underlined. Modified residue is preceded by “d” to signify dehydrated T. ▼ indicates [M+2H-H_2_O]^+2^ ions.

## DISCUSSION

When co-expressed with SEC in *E. coli*, GI is O-GlcNAc modified. Through a combination of deletion, mutational and MS analysis a single amino acid, T829, was shown to be modified. Our analysis suggests that this is the only modified site. We have previously identified several proteins including RGA, TCP proteins and the coat protein of plum pox virus that are also modified in the *E. coli* co-expression system as well as plants (Kim et al., 2011; Scott et al., 2006; Steiner et al., 2012) suggesting that GI is also O-GlcNAc modified in plants. Although O-GlcNAc modification of GI was not identified in several recently reported global analyses of proteins modified in plants (Li et al., 2023; Wu et al., 2022; Xu et al., 2017), GI protein abundance is subject to circadian fluctuations, being most abundant in evening (Krahmer et al., 2019) and thus could have been missed in these global analyses.

Studies identifying O-fucose and O-GlcNAc modified proteins have found that proteins can have both modifications (Bi et al., 2023; Zentella et al., 2023). GI was shown to interact physically and genetically with SPY (Tseng et al., 2004), the Arabidopsis O-fucose transferase (Zentella et al., 2017). It will be interesting to see if SPY also modifies GI and how O-fucose and O-GlcNAc affect GI function.

Three different strategies for enrichment and preparation of modified peptides for MS analysis were tested. All three strategies successfully enriched modified peptides and mapped the modification. Proteins can be modified with N- or O-GlcNAc and strategies that enrich GlcNAc modified peptides and MS do not discriminate between these modifications. β-elimination selectively removes O-linked modifications and dehydrates the modified amino acid. CID MS/MS analysis of purified O-GlcNAc peptides after β-elimination detected dehydration of only T829 suggesting that β-elimination can be used in MS mapping strategies to distinguish between N- and O-linked modifications on purified GlcNAc-modified peptides. A drawback with this method is that other O-linked modifications can be β-eliminated. Therefore, although the peptides are enriched by virtue of being GlcNAc modified it is possible that the peptides will have other O-linked modifications such as O-fucose or phosphorylation. Pretreatment with phosphatase and removal of O-fucose modified peptide using *Aleuria aurantia* lectin should help distinguish between the different modifications.

## AUTHOR CONTRIBUTIONS

Young-Cheon Kim, Lynn Hartweck and Neil Olszewski designed the experiments. Young-Cheon Kim and Lynn Hartweck performed the experiments. Young-Cheon Kim, Lynn M. Hartweck and Neil Olszewski analyzed the data, and wrote and edited the manuscript.

## ACKNOWLEDGMENTS

This work was supported by NSF grants (MCB-0112826, MCB-0820666 and MCB-1158089) to NEO and DOE grant (DE-FG01-04ER04) to NEO and LMH. We thank Matthew D. Stone and LeeAnn Higgins of the University of Minnesota Center for Metabolomics and Proteomics Facility for performing the CID and ETD analysis and assistance with analysis of the results. We thank George Coupland for providing a full-length GI cDNA clone.

---

**Supplemental Table 1.**
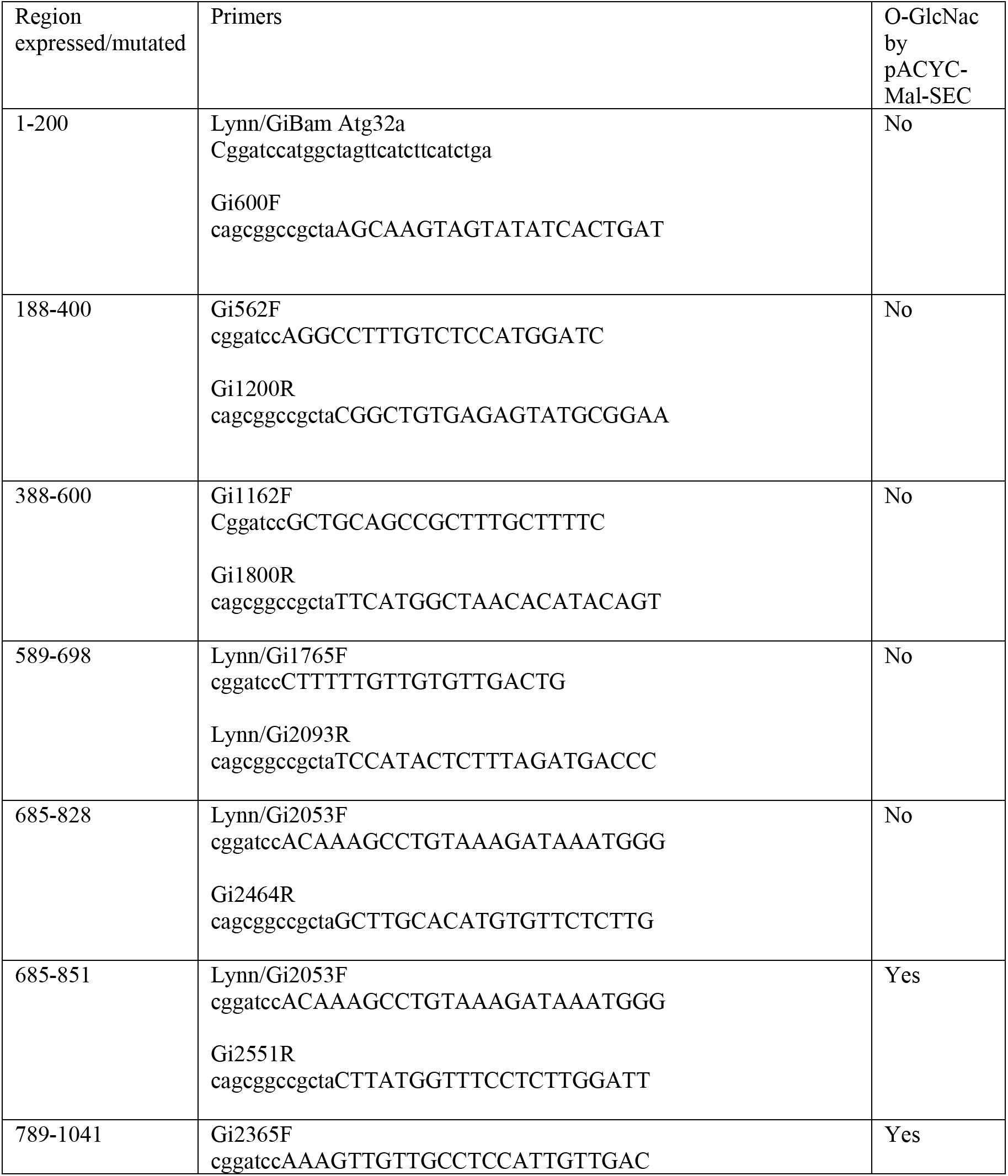
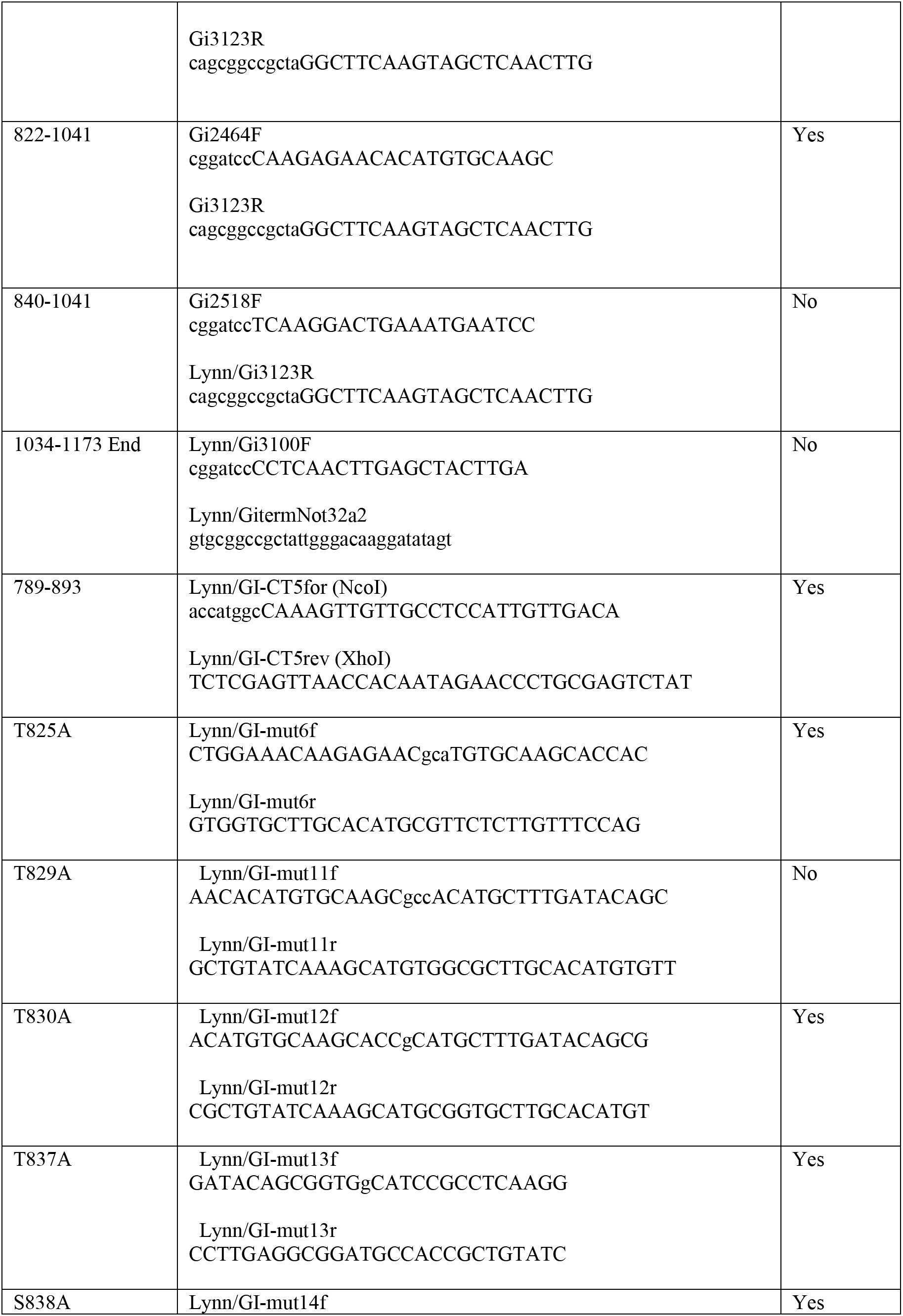
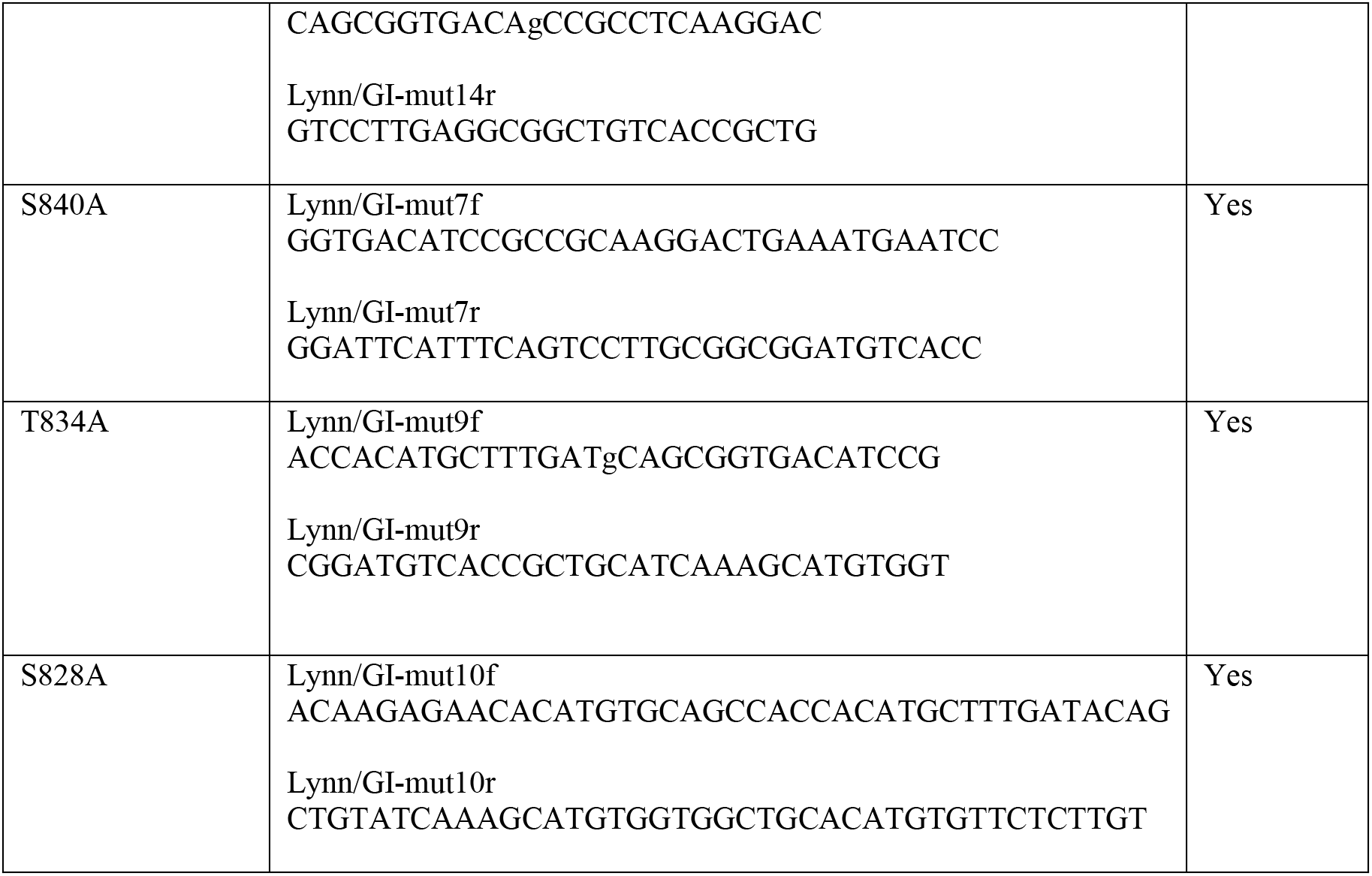
Testing of GI modification across different regions and amino acids of GI. Columns list the region of GI amplified in amino acids. In some cases, substitutions are listed. The forward and reverse primers used to make the GI fragments by PCR amplification are listed. The results of testing whether SEC modified the fragments (shown in Figure 1) are summarized in the last column.

**Supplemental Figure 1.**
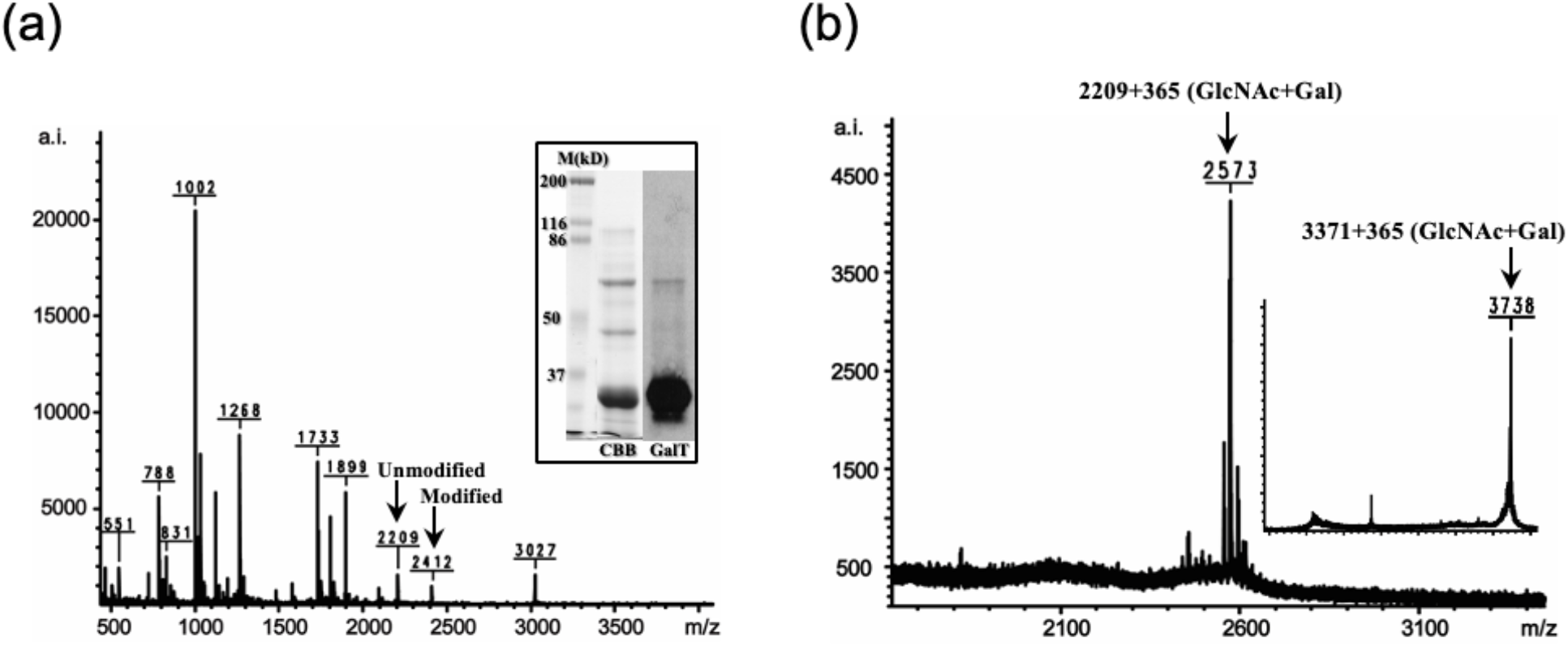
O-GlcNAc modified CT5 peptides were enriched by RCA I affinity chromatography. (a) MALDI-TOF analysis of trypsinized CT5 before capping with galactose and enrichment. Unmodified and modified QENTCASTTCFDTAVTSASR peptides with single O-GlcNAc (203 Da) were observed at *m/z* 2209 and 2412, respectively. The insert shows that CT5 co-expressed with SEC in *E. coli* was O-GlcNAc modified. The panel on the left shows the SDS-PAGE resolving with marker (M) followed by Coomassie Brilliant blue (CBB) and the right panel (GalT) shows the same blot after labeling of GlcNAc residues with [^3^H] galactose. (b) Peptides enriched by RCA I column chromatography was analyzed using MALDI-TOF. After trypsin digestion and enrichment, CT5 QENTCASTTCFDTAVTSASR peptides (*m/z* 2209) containing single modification +365 Da (LacNAc) was observed at *m/z* 2573. The insert shows that QENTCASTTCFDTAVTSASRTEMNPRGNHK peptides (*m/z* 3371) containing a single modification (+365 Da) was observed at *m/z* 3736 after Lys-C digestion and enrichment.

**Supplemental Figure 2.**
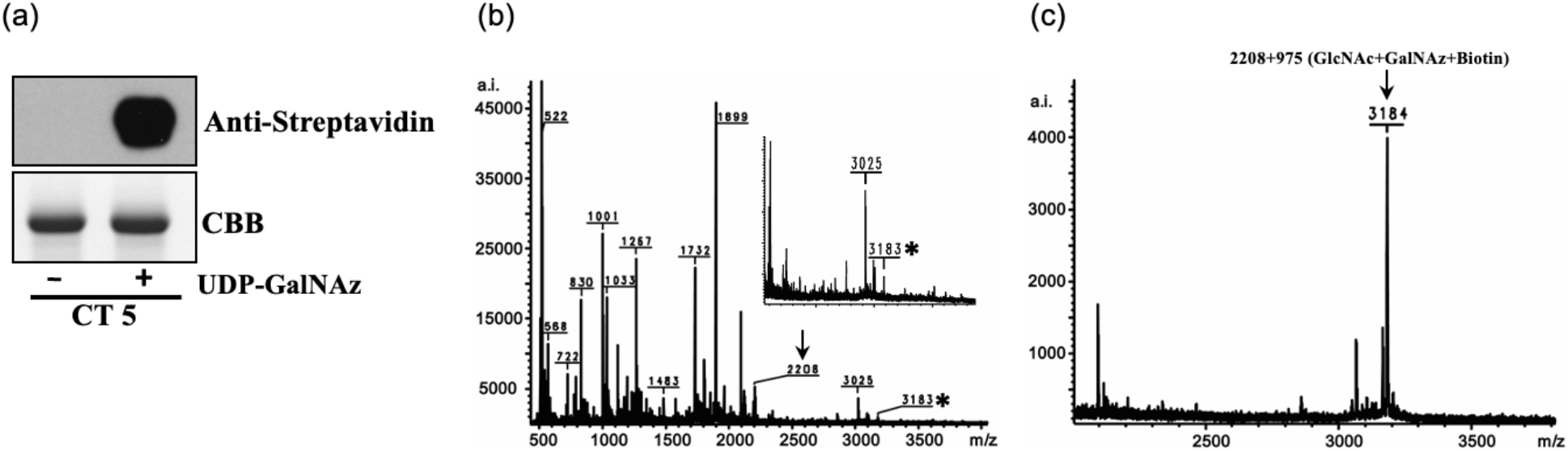
O-GlcNAc modified CT5 proteins were enriched by click enzymatic/chemical labeling and avidin gel purification. (a) The upper blot shows the modified CT5 were labeled with GalNAz and bitoin by click enzymatic/chemical reaction. The lower blot stained with Coomassie Brilliant blue (CBB) shows the amount of CT5 that was loaded in each lane. (b) MALDI-TOF analysis of trypsinized CT5 after click enzymatic/chemical labeling with GalNAz and biotin. The arrow indicates the unmodified QENTCASTTCFDTAVTSASR peptide (*m/z* 2208) and asterisk indicates the peptide modified with GlcNAc-GalNAz-biotin (+975 Da). The insert is an enlargement of a portion of the spectrum. (c) MALDI-TOF results shows that modified was enriched by avidin gel.

**Supplemental Figure 3.**
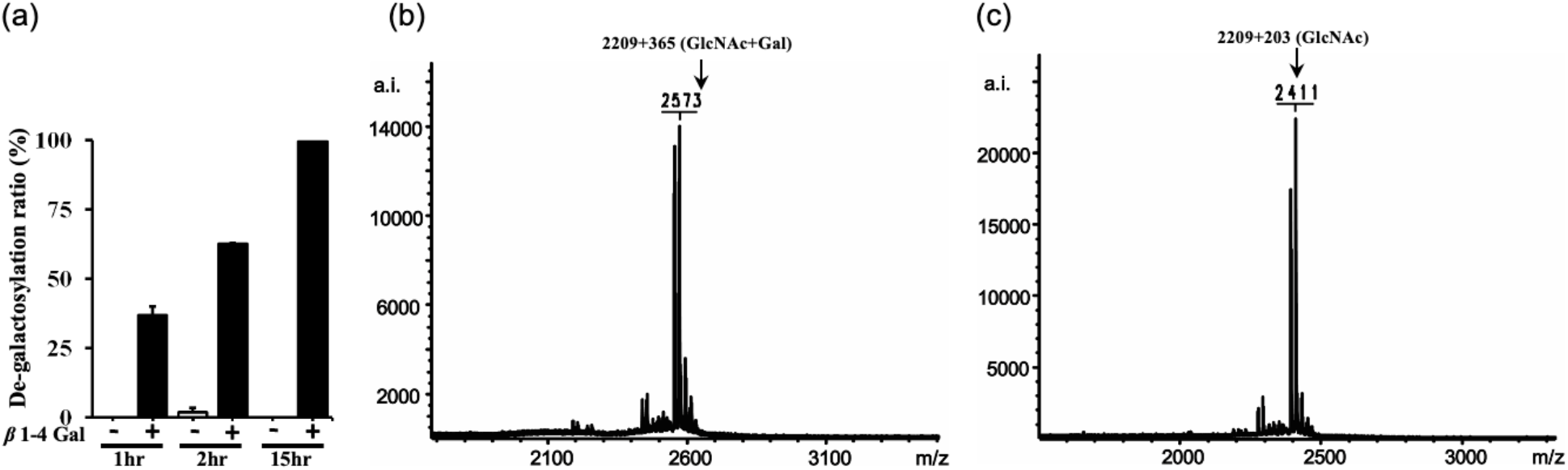
β1-4 Galactosidase cleavage of galactose residues bound to O-GlcNAc. (a) Time course for release of [^3^H]galactose by β1-4 Galactosidase (β1-4 Gal). (b) MALDI-TOF analysis result shows that there is no change the LacNAc moiety after 15hr reaction without enzyme. (c) MALDI-TOF analysis result shows that the β1-4 galactosidase specifically removed galactose.

